# Rapid spatial learning controls instinctive defensive behavior in mice

**DOI:** 10.1101/116236

**Authors:** Ruben Vale, Dominic A. Evans, Tiago Branco

## Abstract

Instinctive defensive behaviors are essential for animal survival. Across the animal kingdom there are sensory stimuli that innately represent threat and trigger stereotyped behaviors such as escape or freezing [1-4]. While innate behaviors are considered to be hard-wired stimulus-responses [5], they act within dynamic environments, and factors such as the properties of the threat [6-9] and its perceived intensity [1, 10, 11], access to food sources [12-14] or expectations from past experience [15, 16], have been shown to influence defensive behaviors, suggesting that their expression can be modulated. However, despite recent work [2, 4, 17-21], little is known about how flexible mouse innate defensive behaviors are, and how quickly they can be modified by experience. To address this, we have investigated the dependence of escape behavior on learned knowledge about the spatial environment, and how the behavior is updated when the environment changes acutely. Using behavioral assays with innately threatening visual and auditory stimuli, we show that the primary goal of escape in mice is to reach a previously memorized shelter location. Memory of the escape target can be formed in a single shelter visit lasting less than 20 seconds, and changes in the spatial environment lead to a rapid update of the defensive action, including changing the defensive strategy from escape to freezing. Our results show that while there are innate links between specific sensory features and defensive behavior, instinctive defensive actions are surprisingly flexible and can be rapidly updated by experience to adapt to changing spatial environments.

## Results

### Escape behavior is a goal-directed action to reach safety

When escaping from imminent threat, animals have two general options: to move away from the threat or to move towards safety. These two behaviors have different consequences and are fundamentally distinct in the computations they require. Moving away from threat can be implemented as a simple reaction to the stimulus [22], but it has the drawback that it might not be the most adaptive solution, if it increases detectability or the animal moves into a position where it cannot escape from [3, 23]. On the other hand, moving towards a safe place has better long term value, but requires more complex computations that might take valuable time, such as evaluating shelter locations and available escape routes. To test which strategy is preferentially used by mice exposed to innately aversive threats, we placed naïve animals in a Barnes maze, which is a circular arena with 20 identical holes that are all covered except for one that leads to an underground shelter ([24], Figure 1A). After a short habituation period (7min) during which mice spontaneously found the shelter location, we exposed them to overhead dark expanding spots, previously shown to be innately aversive [4], delivered either in-between the mouse and the shelter (on-path), or directly above the mouse (on-top). Both stimuli elicited fast escape to the shelter with short reaction times (202±16ms, n=51 responses from 26 animals; Figure 1B; Movie S1) independently of the initial location of the mouse (Figure 1C). Surprisingly, we found no relationship between the stimulus position and the evoked escape trajectories, which were all directed to the shelter, even when the stimulus was between the mouse and the shelter, requiring the mouse to run towards the aversive stimulus in order to reach safety (Figure 1B). In contrast with trajectories during foraging, flight trajectories were very close to a straight line, and not different between the two stimulus conditions, (mean linearity ratio: on-path 106±1%, on-top 109±2%, foraging 209±30%; p=0.27 *t-test* between on-path and on-top, p<0.0001 *t-test* between flights and foraging), as well as highly accurate (mean accuracy: on-path 89±5%, on-top 97±1%, p=0.32 *t-test* between on-path and on-top) despite the lack of any long-term training (Figure 1D-E). In addition, the first body movement after the onset of the stimulus was head orientation towards the shelter location. This orienting behavior was independent of the initial angle between the head direction and the shelter, which was reduced to less than 10 degrees before the mouse covered the first 10% of the distance to shelter, and thus preceded the onset of full flight (Figure 1F-G and Figure S1). Remarkably, in 91.5% of the trials mice rotated their head towards the side of the narrower angle, indicating an awareness of the flight target before the onset of head turning. Similar behavior was observed in response to overhead ultrasonic sweeps [25], which represent a more spatially diffuse threat (Figure 1B-G; see Experimental Procedures), and further support the independence of the behavior from threat localization in this environment.

**Figure 1.**
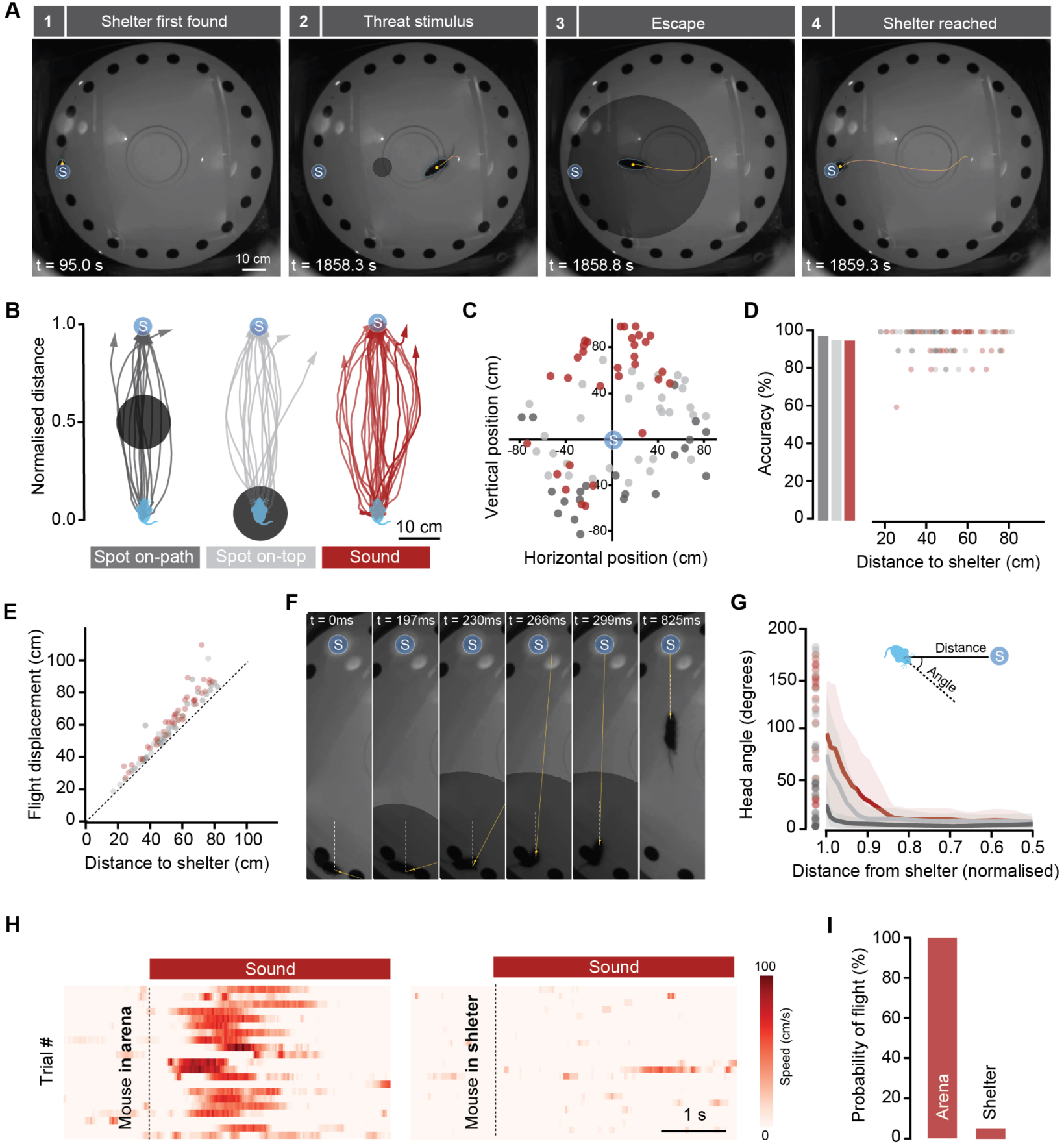
Escape behaviour is a goal-directed action to reach safety. **(A)** Video frames from one trial showing escape to a previously explored shelter after stimulation with an expanding spot projected from above, between the mouse and the shelter location (on-path). Yellow lines indicate the mouse trajectory during the preceding 2 sec. **(B)** Example trajectories from several mice, recorded between stimulus onset and the end of flight, showing that flight path and target are independent of stimulus location or quality. **(C)** Initial position of mice in all trials plotted in relation to the shelter location. **(D)** Accuracy of reaching the shelter during escape. Bars show average accuracy and circles are individual accuracy data points as function of distance to the shelter. **(E)** Total displacement during escape for **100%** accurate flights plotted against linear distance to the shelter. **(F)** Video frames from one trial during initiation of escape from an expanding spot on-top, highlighting the initial head rotation preceding the initiation of running. Yellow line indicates head direction and dashed white line is the reference line between the current mouse position and the shelter. **(G)** Head angles measured between the white and yellow lines illustrated in **(F)** for **100%** accurate flights, showing that the head is pointing towards the position of the shelter before the distance to the shelter is covered. Circles are the initial angles for different trials, lines are average head rotation profile and shaded areas represent SD. **(H)** Raster plots showing speed profile of trials in several mice stimulated with sound when exploring the arena (left) or when the same mice where inside an overground shelter (right). **(I)** The probability of flight is dramatically reduced when animals are already inside a shelter. For all relevant panels: blue circle with “S” identifies the shelter location, dark gray, light gray and red colours represent data from stimulation with spot on-path, spot on-top and sound, respectively.

These data suggest that the goal of the escape behavior is to reach safety. To further test this hypothesis, we reasoned that presentation of the threat while the animal is in the shelter should not cause escape behavior. Indeed, auditory stimuli delivered both in the Barnes maze and in a modified version with a surface shelter, did not cause escape behavior despite the sound pressure level inside the shelter being within 2dB of the arena outside (escape probability = 100% outside vs 6% inside, p<0.001, *t-test* between the two conditions, n=76 responses from 11 animals; Figure 1H-I), indicating that the perception of safety can veto escape from innately aversive threats. These results show that instinctive escape behavior in the mouse is not a simple stimulus-reaction, but a generic action in response to threat with the goal of reaching a safe area, the location of which is computed before the onset of the escape.

### Memory of shelter location guides defensive flight

We next investigated the strategies mice use to determine shelter location. Previous work has shown that foraging rodents can navigate using a variety of strategies [26, 27], including retrieval of a cognitive spatial map [28], relying on prominent external landmarks [26] or integrating self-motion cues over time (path integration, [29, 30]). Here, we tested whether spatial landmarks in the local surroundings of the shelter are used to guide escape, and whether flight termination is signaled by the safety conferred by arriving inside the shelter. We performed two complementary experiments. First, we placed animals in a modified Barnes maze where the center is fixed and the periphery can be automatically rotated, together with a set of olfactory and visual local cues that have been shown to guide navigation in mice [31]. Escape responses to the shelter were first elicited with sound stimuli, after which the peripheral ring of the arena was rotated by a random angle when mice were in the center (range: 36°-90°, mean=56°; corresponding to 2-5 holes, mean=3.1), and the sound stimulus delivered again (Figure 2A, Movie S2). All mice invariably ran towards the previous shelter location, with accuracy, trajectory linearity, reaction times and head orientation profile that were not different from pre-rotation flights (Figure 2B-D). Moreover, mice stayed in the vicinity of the pre-rotation location for 4.6±0.2s, which is 2.5 times longer than the time mice spent in the wrong location during missed flights in control conditions (Figure 2E, p<0.001 *t-test* for time in the wrong location between control and post-rotation), further indicating goal directedness towards this location. These data suggest that landmarks proximal to the shelter are not required for the computation of shelter location, and this is further supported by threat presentation in complete darkness, which evokes perfectly accurate escape responses (Figure S2A-B, Movie S2). Next, we placed a shelter in the center of the arena, to which mice fled reliably when exposed to auditory stimulation, and then removed the shelter and repeated the auditory stimulation. Remarkably, this resulted in flights that stopped in the arena center (Figure 2F-H, Movie S2) and were followed by persistence in this location, which is normally aversive to mice (Figure S2C), sometimes up to 15 seconds (mean=2.5±1.1s). Together with the previous experiment, these results show that mice escape towards a previously memorized shelter location, and that flight termination is signaled by having reached the stored target location, and does not require reaching safety.

**Figure 2.**
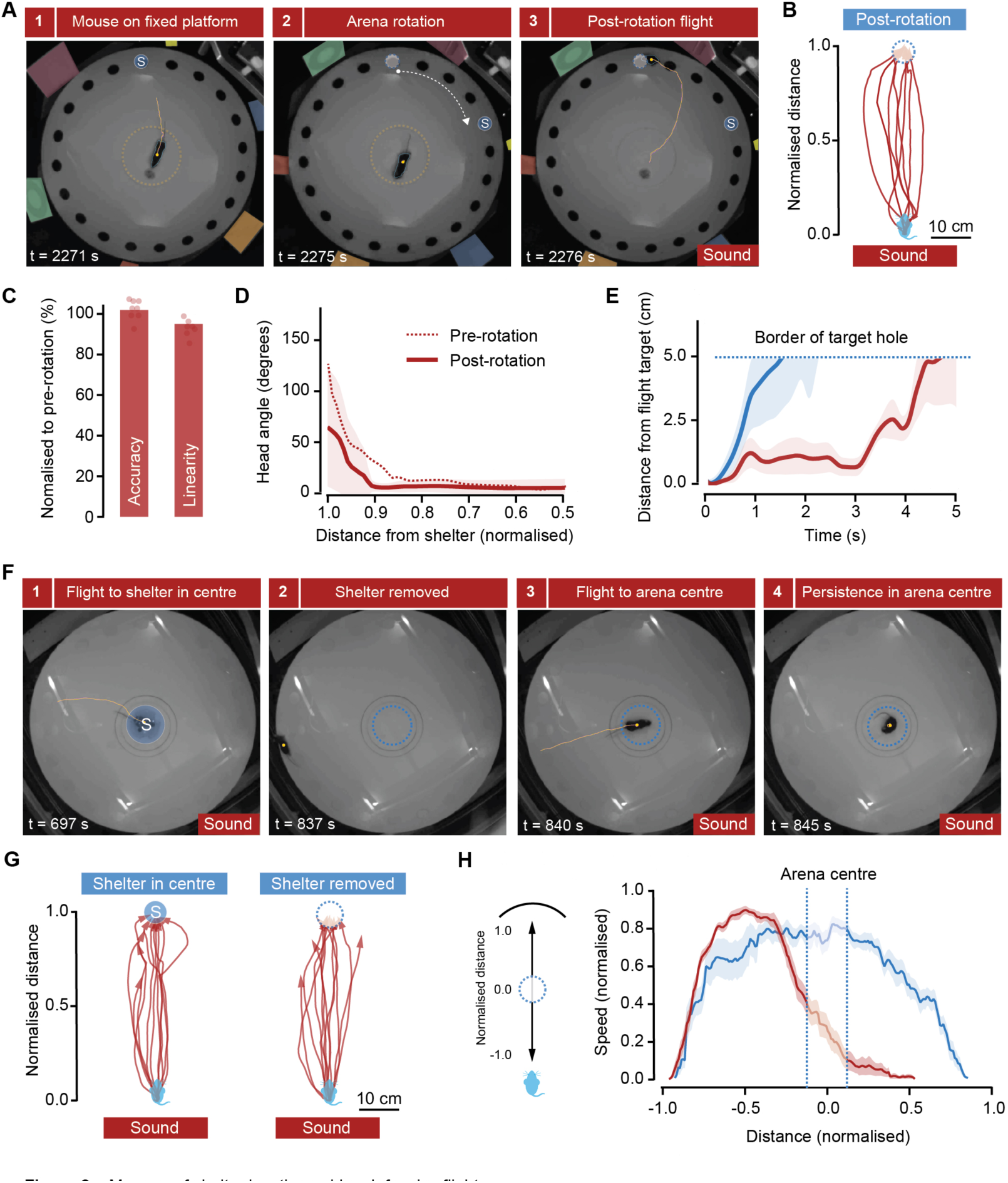
Memory of shelter location guides defensive flight. **(A)** Video frames from one trial showing escape from aversive sound immediately after the outside of the arena had been rotated, together with local cues (panels on the outside, color-coded for clarity). Dashed yellow line marks the diameter of the fixed platform, and dashed blue circle shows shelter location before rotation. **(B)** Trajectories from different mice after arena rotation, showing escape towards the previous shelter location (dashed blue circle). **(C)** Escape behaviour is not significantly changed by arena rotation. **(D)** Head rotation profile during escape initiation is not affected by arena rotation. Post-rotation angles are measured between the mouse position and the shelter position before rotation. Shaded area shows SD. **(E)** Plot showing when the mouse leaves the initial target hole after the flight. Red is for flights after rotation and blue is for flights in control conditions where the shelter target was missed. Shaded area shows SEM. **(F)** Video frames from one trial showing sound-evoked flight to a shelter in the centre of the arena, and persistence of escape to the arena centre after the shelter has been removed. **(G)** Escape trajectories for different mice before (left) and after (right) a shelter in the arena centre was removed. **(H)** Speed profile for escape responses when the shelter is in the periphery (blue, from the same dataset shown in Figure 1) and after the shelter has been removed from the arena centre.

### Shelter location memory is formed rapidly and supports fast updates in defensive actions

If mice rely on memory of the shelter location to reach it, how is this memory formed? To determine this, we removed the fixed habituation period and exposed animals to threat immediately after they visited the shelter for the first time. Even though animals were inside the shelter for as little as 18 seconds (range: 18-270s, n=12 animals; Figure 3A), this was enough to support shelter-directed escape responses that were indistinguishable from the control condition (Figure 3B; p=0.79 for accuracy and p=0.78 for linearity, *t-test* against control). This shows that memory of shelter location is formed by a very fast single-trial learning process. Interestingly, there was a significant negative correlation between the total time spent in shelter and the reaction time (Pearson’s r=-0.46, p=0.007), suggesting that computation of the escape vector might depend on the strength of the shelter location memory (Figure 3C).

We next investigated how shelter place memory supports updates in defensive actions when the environment changes, by performing two sets of experiments. First, we elicited one flight with the sound stimulus in control conditions, after which we changed the location of the shelter to the opposite hole (see Supplemental Experimental Procedures). We then waited until animals spontaneously visited the new shelter (mean time 33.1s, range: 4-82s), after which we ran several trials of sound stimulation. We found that animals escaped to the new shelter location in less than two trials (mean=1.8±0.3 trials), with 4/9 of mice escaping to the new location on the first trial. Some animals still escaped to the old location after having fled to the new one on a previous trial, but after 4 trials (mean value; over a period of 10.5±6.8min), 9/9 of animals escaped repeatedly to the new location (Figure 3D-E). Importantly, escapes to the old location were immediately followed by secondary straight flights to the new location (including 4/5 first trial escapes to the old location, Figure 3D, Movie S3), suggesting that despite reaching the wrong target mice already hold the memory of the new shelter location. This shows that the new shelter location can be stored in a single trial, and that safety devaluation of the old location supports a permanent update of the escape target after a small number of trials. In the second set of experiments, we closed the shelter hole, and after 7min of exploration, during which animals always visited the closed shelter location, presentation of the looming stimulus directly above the mouse did not elicit escape, but instead caused freezing for the duration of the stimulus (freezing probability = 71.4%, mean freezing time = 629.9±100.0ms; flight probability = 10.7%; Figure S3), including long lasting freezing for slowly expanding spots, sometimes lasting a long as 50s (freezing probability = 95.2%, mean freezing time = 7.9±2.7s; flight probability = 4.8%; Figure 3F-G, Movie S3). This change in defensive strategy was completely reversible, as stimulus presentation 5min after re-opening the shelter hole once again produced robust shelter-directed flights (Figure 3G, Figure S3). These data show that instinctive defensive escape is conditional on the knowledge of an existing shelter location, and that in the absence of a memory of shelter location, mice switch their defensive strategy to freezing.

**Figure 3.**
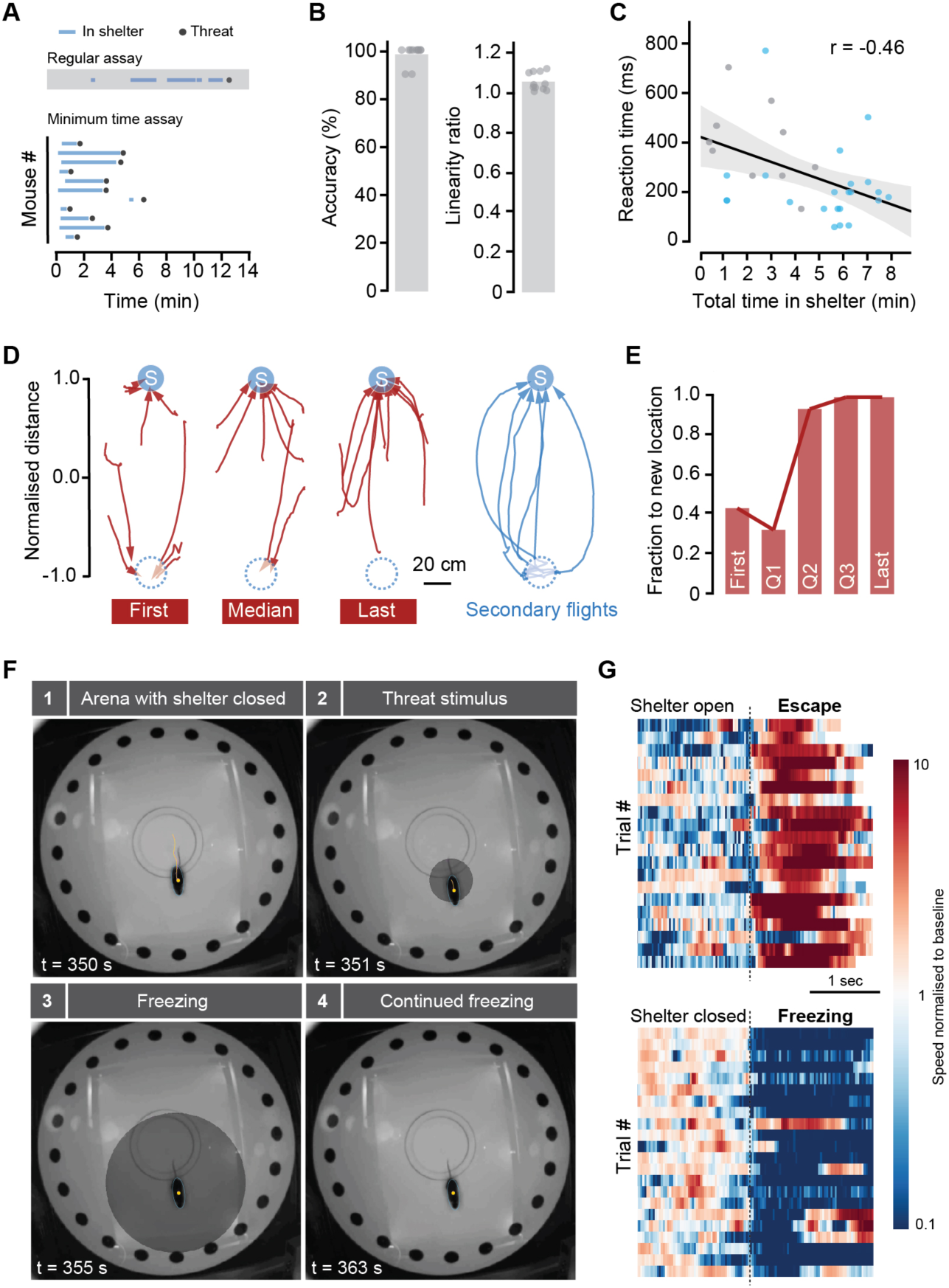
Shelter location memory is formed rapidly and supports fast updates of defensive actions. **(A)** Raster plot showing periods of time inside the shelter from the onset of arena exploration, and threat stimulus presentation. Top, example raster from a regular assay for comparison (as shown in Figure 1), with multiple entries in the shelter during the exploration phase. **(B)** Average (bars) and data points (circles) for accuracy and linearity of escape after shelter single visits. **(C)** Time to initiate escape is negatively correlated with the total amount of time spent in the shelter before stimulation. Gray circles are data from the minimum time assay, and blue circles are data from the regular assay. Black line is a regression line fit to all data points, shaded area is 95% confidence interval for the regression. **(D)** Escape trajectories after the original shelter has been closed (dashed blue circle), and a new one open in a different position (blue circle with “S”), for the first and last trials (left and right red, respectively), and the median trial (centre red). Trajecories in blue (right) are for secondary flights, which immediately follow escapes to the original location.**(E)** Evolution of escape behaviour after shelter location has been moved, as in **(D)**, showing the fraction of flights across all mice that reach the new shelter location, for first, three quartiles (Q1-Q3), and last trials. **(F)** Video frames from one mouse in an arena with the shelter closed, showing freezing behaviour in response to a slowing expanding spot projected on top. **(G)** Raster plots showing speed profiles upon threat stimulation before (bottom) and after the shelter hole has been opened (top), for slowly expanding spots. Trials have been aligned by reaction time (dashed line).

## Discussion

We have shown that instinctive defensive actions depend on rapidly learned information about the spatial environment, and that the expectation of safety drives escape behavior to a learned shelter location, while its absence promotes defensive freezing. Our results support the idea that computations other than threat detection play an important role in the initiation of defensive behavior [32]. In our assay, there are at least two computational steps that precede defensive action: evaluation of whether shelter is available, and if so, determination of its location. The first is used to choose between fleeing or freezing, and the second to compute an escape vector from the current position to the shelter location, which we demonstrate to happen before flight initiation. Importantly, we show that information about the availability and location of the shelter is stored as a memory, and therefore suggests that mice use spatial representations to coordinate instinctive defensive behaviors. This is in agreement with results from experiments in gerbils suggesting that spatial maps might be used to optimize escape routes [33]. In our experiments, the same visual stimulus could elicit both escape and freezing depending on the spatial configuration of the arena, and thus while it is possible that different defensive behaviors might be mediated by distinct visual pathways, as previously suggested [2], our results are compatible with a more general model where sensory stimuli are incorporated into higher-order information streams to make the choice between freezing and fleeing from threat.

Previous studies on foraging rodents have shown that spatial navigation can be accomplished using both landmark information and self-motion cues, and that when both are present the most reliable information is used [29, 34, 35]. For example, homing hamsters will follow local cues that have been rotated, but only up to a certain angle, after which they switch strategies and perform path integration [34]. In our experiments, rotation of local landmarks did not change the accuracy of escape behavior, suggesting that self-motion cues might play an important role when fleeing from threats. While we cannot rule out that landmarks outside our experimental control contribute to navigation, path integration is particularly well suited to compute the current position as a vector from a home base [36], and could be the preferred strategy during escape. This strategy might have the advantage that animals do not need to scan the environment for local cues that signal the shelter, which could take a significant amount of time, and might thus shorten computation times. Interestingly, mice stop at the learned shelter location when the shelter is absent, even if the location is the arena center, suggesting that shelter cues and the safety conferred by the shelter are not processed during the escape response, and are not necessary to terminate flight. A key finding of this study is that learning the shelter location is a very fast process requiring only a single visit, and that flight accuracy is extremely high from the first escape trial. This contrasts with previous experiments using Barnes mazes, where the accuracy to find the shelter increases slowly over multiple trials across several days [31, 37]. An important difference is that, in our experiments, threats were presented after mice moved away from the shelter voluntarily instead of being placed in the maze center by the experimenter [31, 37], further supporting the idea that path integration might be the dominant navigation strategy during escape.

A key consequence of rapid spatial learning is that it greatly increases the flexibility of escape behavior. We have shown that a single, short-lived visit to a shelter is sufficient to support accurate escape behavior, and that changes in the environment are incorporated into action selection within minutes, suggesting that mice have very rapid mechanisms for risk assessment [38, 39]. Importantly, when we devaluated the outcome of the flight by moving the shelter to a new location, mice updated the flight goal within a few trials, indicating that the expected outcome of the defensive action might be taken into account, and that instinctive escape could be considered within a model-based behavior framework [40]. In conclusion, while instinctive defensive behaviors rely on innate stimulus-response associations, their computation takes into account internal models of the world that are rapidly updated, and we suggest that they are a powerful model for investigating the neural basis of motivated action selection.

## Experimental Procedures

### Animals and Behavioral Procedures

Male C57BL/6J mice were used for experiments at 6-12 weeks old and tested during the light phase of the light cycle. The main behavioral arena used was a modified Barnes maze [24], consisting of a white acrylic circular platform 92cm in diameter with 20 equidistant circular holes The central area of the arena was a fixed circular platform, and the periphery was mounted on a frame that allowed rotation over 360 degrees. The maze was surrounded by visual cues and bedding from the home-cage of the mouse being tested was placed inside the shelter. Experiments were recorded at 30-50 frames-per-second with a near-infrared camera. Unless otherwise noted, animals were given a 7min acclimation period, and an additional 5min if they did not visit the shelter at least once. If the shelter was not found in this period, the experiment was terminated.

### Auditory and Visual stimulation

The auditory stimulus consisted of a train of three frequency modulated upsweeps from 17 to 20kHz over 3s [25], lasting 9s in total, at a sound pressure level of 73-78dB as measured at the arena floor. Visual stimuli were backprojected on to a screen positioned 64cm above the arena and consisted of an expanding dark circle (Weber contrast, -0.98) on a gray background (luminance, 7.95cd/m^2^) [4]. The standard circle subtended a visual angle of 2.6° at onset and expanded linearly at 224°/s over 200ms to 47.4°, at which it remained for 250ms. In Figure 3G, the expansion rate of the circle was 11.2°/s over 4s, and the expanded size was maintained for 1250ms.

## Author Contributions

R.V. and T.B. designed the study and experiments. R.V. performed all experiments with assistance from D.E. R.V and T.B. analyzed the data. T.B. wrote the manuscript with input from R.V. and D.E.

## Acknowledgments

This work was funded by a Wellcome Trust/Royal Society Henry Dale Fellowship (098400/Z/12/Z), a Medical Research Council (MRC) grant MC-UP-1201/1, a Wellcome Trust and Gatsby Charitable Foundation SWC Fellowship (to T.B.), MRC PhD Studentship (to D.E. and R.V) and a Boehringer Ingelheim Fonds PhD fellowship (to R.V). We thank Kostas Betsios for programming the data acquisition software, the LMB Mechanical and Electrical Workshops for building the experimental arenas, P. Dayan, T. Mrsic-Flogel, C. Schmidt-Hieber and members of the Branco lab for discussions, and S. Sternson, J. O’Keefe, P. Dayan, K. Lloyd, T. Margrie and I. Bianco for comments on the manuscript.

---

**Movie S1** – Examples of mice spontaneously finding the shelter during exploration, and fleeing to the shelter in response to expanding spots delivered on-path and on-top, and to ultrasonic sweeps.

**Movie S2** – Illustrative defensive responses to threat after acute spatial changes in the arena, and when there is no arena illumination.

**Movie S3** – Example of a secondary flight after switching the shelter location, and of a freezing response to a slowly expanding spot in the absence of shelter.

